# *N*-glycosylation on *Oryza Sativa* Root Germin-like Protein 1 is conserved but not required for stability or activity

**DOI:** 10.1101/2021.02.09.430526

**Authors:** Tehseen Rubbab, Cassandra L. Pegg, Toan K. Phung, Amanda S. Nouwens, K. Y. Benjamin Yeo, Lucia F. Zacchi, Amna Muhammad, S. M. S Saqlan Naqvi, Benjamin L. Schulz

## Abstract

Germin and germin-like proteins (GLPs) are a broad family of extracellular glycoproteins ubiquitously distributed in plants. Overexpression of *Oryza sativa* root germin like protein 1 (*Os*RGLP1) enhances superoxide dismutase (SOD) activity in transgenic plants. Here, we report bioinformatic analysis and heterologous expression of *Os*RGLP1 to study the role of glycosylation on *Os*RGLP1 protein stability and activity. Sequence analysis of *Os*RGLP1 homologs identified diverse *N*-glycosylation sequons, one of which was highly conserved. We therefore expressed *Os*RGLP1 in glycosylation-competent *Saccharomyces cerevisiae* as a Maltose Binding Protein (MBP) fusion. Mass spectrometry analysis of purified *Os*RGLP1 showed it was expressed by *S. cerevisiae* in both *N*-glycosylated and unmodified forms. Glycoprotein thermal profiling showed little difference in the thermal stability of the glycosylated and unmodified protein forms. Circular Dichroism spectroscopy of MBP-*Os*RGLP1 and a N-Q glycosylation-deficient variant showed that both glycosylated and unmodified MBP-*Os*RGLP1 had similar secondary structure, and both forms had equivalent SOD activity. Together, we concluded that glycosylation was not critical for *Os*RGLP1 protein stability or activity, and it could therefore likely be produced in *Escherichia coli* without glycosylation. Indeed, we found that *Os*RGLP1 could be efficiently expressed and purified from K12 shuffle *E. coli* with a specific activity of 1251±70 Units/mg. In conclusion, we find that some highly conserved *N*-glycosylation sites are not necessarily required for protein stability or activity, and describe a suitable method for production of *Os*RGLP1 which paves the way for further characterization and use of this protein.

## Introduction

Germins and germin-like proteins (GLPs) belong to a functionally diverse family of proteins called cupins [1]. These proteins have been extensively studied in cereals and legumes due to their historical significance, abundance, expression during biotic and abiotic stresses and potential biotechnological applications [2, 3]. Germins were first identified during a search for molecular markers associated with germination in wheat and were the only gene product synthesized *de novo* during germination; hence the name germin, implying an association with and serving as a signal for germination [4]. Several germin isoforms have been identified in diverse cereals, but not all are associated with germination. GLPs form a broader protein family, with ~30-70% sequence similarity to wheat germins [5].

Germins and GLPs share a well conserved β-barrel fold [1] and are typically glycoproteins. Differences in the extent of glycosylation are responsible for their presence as different isoforms, with variably modified forms associated with different subcellular localization [6, 7] and tissue distribution [8]. *Oryza sativa* root germin like protein 1 (*Os*RGLP1) contains a putative *N*-glycosylation site (N-x-S/T; x≠P) [9], but the exact role of glycosylation in *Os*RGLP1 protein stability, folding, or activity is unknown. Glycosylation affects the folding efficiency and final structure of proteins, and most secreted proteins are glycosylated. Changes in glycosylation can affect how glycoproteins recruit, interact with, and activate other proteins in diverse biological systems [10]. In addition to assisting in protein folding, glycosylation is also important for increasing protein thermostability. Addition of even a single glycosylation event can impact the equilibrium of protein between folded and unfolded states, although in some proteins elimination of all or some glycans does not affect protein folding [11].

Germins and GLPs have been widely exploited in transgenic plants to enhance resilience to biotic and abiotic stresses. The most important biochemical activity reported for true germins is oxalate oxidase activity [12], and for GLPs superoxide dismutase (SOD activity [13]. *Os*RGLP1 has been successfully transformed into *Nicotiana tabacum* (tobacco), *Solanum tuberosum* (potato) and *Medicago truncatula* (barrelclover) [13, 14]. *Os*RGLP1-transformed transgenic plants have higher SOD activity than wild type plants, and enhanced stress tolerance due to this heat resistant SOD activity [15]. Heterologous expression of *Os*RGLP1 in potato also induces resistance against infection by *Fusarium oxysporum f.sb. tuberosi* [14]. Transgenic *Medicago truncatula* expressing *Os*RGLP1 are more resistant to *F. oxysporum* than wild type plants, which was attributed to the heat resistant SOD activity of *Os*RGLP1 [16].

Here, we investigated the importance of glycosylation for the stability and activity of *Os*RGLP1. As *Os*RGLP1 likely required eukaryotic post-translational modifications, we first tested the importance of the single conserved glycosylation site in *Os*RGLP1 for protein activity and stability by its expression and purification as an MBP-fusion protein in *Saccharomyces cerevisiae*. Surprisingly, we found that although it contained a conserved *N*-glycosylation site, this was not required for its stability or function. We therefore also produced *Os*RGLP1 in *Escherichia coli*, without glycosylation, and found it was active and could be efficiently expressed and purified, which will be of utility in diverse applications.

## Materials and Methods

### In Silico Analysis of Glycosylation in Germins and GLPs

Homologous sequences to *Os*RGLP1 (UniProtKB accession Q6YZY5) were found using blastP with default parameters. Phylogenetic trees were constructed from alignments by PhyML [17] and ancestral sequence reconstruction was carried with CODEML [18] with a WAG amino acid substitution matrix and molecular clock turned off. The possible instance of glycosylation sites in relation to each other were counted and the intensity of data was displayed as a heat map for visualization of conserved glycosylation sequons.

### Construction of Expression Vectors

DNA encoding Gas1 signal sequence, mature Maltose Binding Protein (MBP) and a factor Xa cleavage site was PCR amplified from vector pRS415-ssGas1-MBP-Gas1-His-Flag [19] with primers ssGas1_SpeI_Fwd (5’-ATCCACTAGTATGTTGTTTAAATCCCTTTC-3’) and MBP_FXa_BamHI_Rvs (5’-ATCCGGATCCCCTTCCCTCGATCCCGAGG-3’) and cloned into pRS425. DNA encoding *Os*RGLP1 without its signal sequence was then PCR amplified from pH7WG2.0 harboring *Os*RGLP1 [14] with primers LZBS66_Fwd (5’-TATAGGATCCGCTTCTGATCCCAGCCCTC-3’) and LZBS70_Rvs (5’-TATACTCGAGTCAGTAATGGTTGTTCTCC-3’) incorporating BamH1 and Xhol sites and cloned into the resulting vector, creating pRS425-MBP-*Os*RGLP1. Site directed mutagenesis was used to mutate *Os*RGLP1 asparagine (Asn/N) at position 76 to glutamine (Gln/Q) using primers LZBS72_Fwd (5’-GTCACACTGATCAACGTCATGCAG-3’) and LZBS73_Rvs (5’-TTGAGACCCAACTTTGTTCGTCTTCCG-3’) as described [20]. DNA encoding mature *Os*RGLP1 was PCR amplified with primers *Os*RGLP1_pET28_Fwd (5’-TATAGGATCCATGGCTTCGTCTTCC-3’) and *Os*RGLP1pET28_Rvs (5’-TATACTCGAGTCAGTAATGTTGTTCTCC-3’) incorporating BamH1 and Xhol restriction sites and cloned into pET-28a(+), resulting in pET-28a(+)-*Os*RGLP1.

### Expression and Protein Purification

*S. cerevisiae* BY4741 (MATa *his3Δ1 leu2Δ0 met15Δ0 ura3Δ0*) was transformed with pRS425-MBP-*Os*RGLP1 or mutant thereof using standard protocols [21]. For protein purification, yeast cells carrying pRS425-MBP-*Os*RGLP1 were grown in SD-Leu media at 30°C with shaking and collected at an OD_600nm_ of 1.0. Cells were resuspended in lysis buffer (50 mM Tris-HCl buffer pH 7.4, 150 mM NaCl, 1x complete protease inhibitor cocktail (PIC) (Roche) and 1 mM PMSF). Cells were disrupted with a OneShot cell disruptor (Constant Systems) at 30 kPsi. Protein was purified using amylose-agarose resin (Clontech) and eluted in 10 mM maltose, 50 mM Tris HCl buffer pH 7.4, and 150 mM NaCl.

K12 shuffle *E. coli* (NEB-c3026) was transformed with pET-28a(+)-*Os*RGLP1 and grown in 100 mL LB media with 100 mg/L kanamycin at 30 °C to an OD_600nm_ 0.6. IPTG (1 mM) was added and incubated at 16 °C, 25 °C or 37 °C shaking at 220 rpm overnight. Cells were collected by centrifugation at 18,000 rcf for 10 min at 4 °C, resuspended in 50 mM Tris-HCl buffer pH 7.4, 150 mM NaCl, 1x PIC, and 1mM PMSF and lysed with a OneShot cell disruptor at 20 kpsi. Protein was purified using Talon resin (Clontech), eluted in 200 mM imidazole, 50 mM Tris-HCl buffer pH 7.4, and 150 mM NaCl, and analyzed by SDS-PAGE. To purify *Os*RGLP1 denatured *E. coli* proteins were prepared as described [22] with slight modifications: cells from 100 mL LB overnight culture of *E. coli* were resuspended in 50 mM Tris-HCl buffer pH 7.4, 150 mM NaCl, 1mM PMSF, 5 mM imidazole, and 1x PIC and lysed with a One Shot cell disruptor at 20 kpsi. The supernatant was heat denatured at 65 °C for 10 min, centrifuged at 12,000 rcf, and aliquots of 0.15 mg/ml protein were frozen. *Os*RGLP1 contaminated with GroL was incubated with 250 μL talon resin for 10 min, pre-equilibrated with one matrix volume of 50 mM Tris-HCl buffer pH 7.4 and 150mM NaCl with gentle shaking at 4°C. Talon resin was washed with denatured *E. coli* proteins (0.1 mg/mL) in 10 mM MgATP, 50 mM Tris-HCl buffer pH 7.4, 150 mM NaCl, 1 mM PMSF, and 5 mM imidazole, and then with 10 mM imidazole, 20 mM MgATP, Tris-HCl buffer pH 7.4, and 150 mM NaCl. *Os*RGLP1 was eluted in 200 mM imidazole, Tris-HCl buffer pH 7.4 and 150 mM NaCl.

### Thermal Stability Analysis

Partially glycosylated MBP-*Os*RGLP1 (5 μg of protein in 100 μL Tris-HCl buffer pH 7.8) was incubated at various temperatures for 10 min, then centrifuged at 21,000 rcf for 10 min to separate aggregated and soluble protein. The supernatant was transferred to a new protein LoBind tube and precipitated by addition of 400 μL methanol/acetone (1:1), incubation at −20°C overnight, and centrifugation at 18,000 rcf for 10 min. The supernatant was discarded and the pellet was resuspended in 100 μL 100 mM ammonium bicarbonate, 10 mM DTT with 0.2 μg of trypsin and incubated at 37°C shaking overnight.

### Mass Spectrometry

For intact protein analysis, 0.5 μg of protein was analysed using a Dionex Ultimate 3000 nanoLC system and Orbitrap Elite (Thermo) mass spectrometer. Proteins were desalted on a C4 PepMap300 pre-column (300 μm x 5mm, 5 μm 300Å) using buffer A (0.1% formic acid) at 30 μL/min for 5 min, a gradient of 10-98% buffer B (80% acetonitrile, 0.1% formic acid) over 5 min, and 98% buffer B for 9 min. The MS was operated in positive ion mode with the Orbitrap analyser set at 120,000 resolution. Source parameters included: ion spray voltage, 2.4 kV; temperature, 275°C; SID=10V; S-lens=70V; summed microscans=3; FT xvacuum=0.1. For MBP-*Os*RGLP1 from *S. cerevisiae*, the ion trap was used: SID=30 V; mass range, 800–2000 *m/z*. For *Os*RGLP1 from K12 shuffle *E. coli*, the mass range was 800–1600 *m/z*. Data was deconvoluted using Thermo Protein Deconvolution software across 900-1600 *m/z* with a minimum of 6-10 adjacent charges, target mass of 25 kDa and charge state range 10-100. Deconvoluted data is reported as uncharged, monoisotopic mass. For tryptic peptide analysis, proteins were denatured, reduced/alkylated, and digested to peptides as previously described [23]. Digested peptides were desalted with C18 ZipTips (Millipore) and measured by LC-ESI-MS/MS using a Prominence nanoLC system (Shimadzu) and a TripleTof 5600 instrument (SCIEX) with a Nanospray III interface as described [24].

### Data Analysis

Peptides and proteins were identified using ProteinPilot v5.0.1 (SCIEX) using a custom database containing the *Os*RGLP1 or MBP-*Os*RGLP1 fusion protein sequences. Settings included: sample type, identification; digestion, trypsin; instrument, TripleTOF 5600; cys alkylation, acrylamide; search effort, thorough. For glycosylation analyses, data was searched with Byonic v2.13.17 (Protein Metrics) against a protein database containing the sequence of the fusion protein and a contaminant protein database. A maximum of two missed cleavages were allowed and mass tolerances of 50 ppm and 75 ppm were applied to precursor and fragment ions, respectively. Cys-S-beta-propionamide was set as a fixed modification and variable modifications were set as mono-oxidised Met and deamidation of Asn. The glycan database contained high mannose glycans (GlcNAc_2_Man_1-15_), with one glycan allowed per peptide.

### Circular Dichroism Spectroscopy

CD spectra were measured from 195 to 280 nm with a Jasco J-710 spectrometer (Jasco) in a 1 mm path length cuvette.

### SOD Assays

SOD assays were performed as described [25]. In-solution SOD activity was determined by the difference between the absorbance of the samples in light and dark. The % inhibition was calculated by the difference between OD_595_ of blank and OD_595_ of sample, divided by OD_595_ of blank. 1 unit of SOD was defined as the amount of SOD required to inhibit the photochemical reduction of NBT by 50% [25]. In-gel SOD activity was measured by separating protein by seminative PAGE gel and negative staining for SOD activity by incubating in the dark in 100 mL 0.1 M potassium phosphate buffer pH 7.8, 0.1mM EDTA, 200 μL TEMED, 20 mg NBT and 30 μL riboflavin with continuous shaking for 30 min. The gel was rinsed twice with distilled water and placed in an illumination chamber for ~15-20 min. The regions where SOD activity was present were negatively stained. *Os*RGLP1 was incubated at various temperatures for 1 h, and remaining SOD activity measured as described above. The data was analyzed in Graphpad Prism 7 using an unpaired *t*-test.

## Results and Discussion

### Conservation of Glycosylation in Germins and Germin-Like Proteins

Germins and GLPs are typically *N*-glycosylated. These post-translational modifications may be important in determining protein structure and activity. Although glycosylation is common in germins and GLPs, their glycosylation site conservation has not been carefully studied. We therefore examined the presence and location of glycosylation sites in proteins with at least 60% sequence homology to *Os*RGLP1. Multiple sequence alignment and phylogenetic analysis revealed that germins and GLPs in rice typically have one or two glycosylation sites. In order to elucidate evolutionary relationships between the different sequon positions we created a correlation heat map (Fig. 1) showing the probability of any two sequons occurring in the same sequence. This revealed a high conservation for N_159_VT (corresponding to N_76_ in *Os*RGLP1), demonstrated by the intense horizontal bar showing conservation of N159 no matter which other glycosylation sites were present, while the presence of glycosylation sequons at other positions was less well conserved. In the rare instances where glycosylation at N_159_ was absent, a sequon was often present at N_150_. Thus, we concluded from this analysis that germins and GLPs in rice have a highly conserved glycosylation site at N_159_, the single glycosylation site present in *Os*RGLP1.

**Figure 1.**
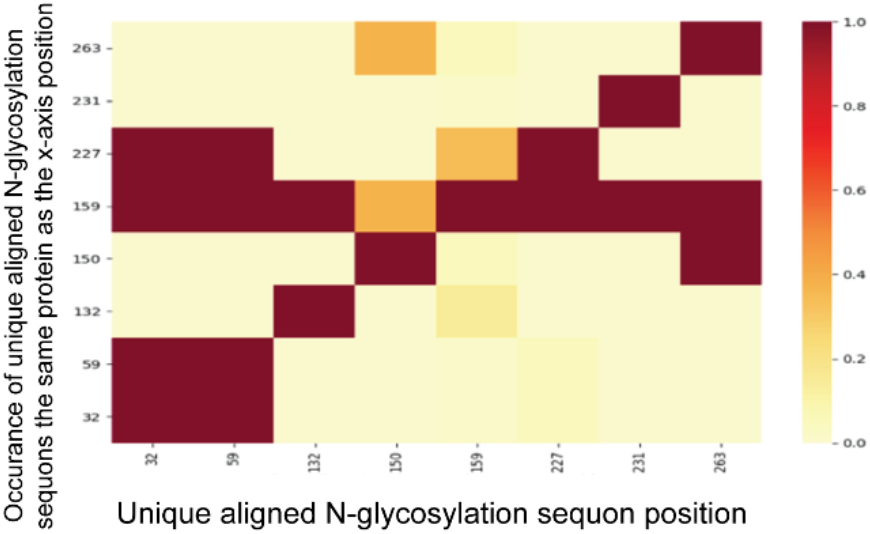
Glycosylation at N_159_ is highly conserved in germins, corresponding to *Os*RGLP1 N_76_. Heatmap cross-correlation of glycosylation sequon conservation in homologs from rice with >60% homology to *Os*RGLP1. X-axis, all unique Asn in sequons. Y-axis, all unique Asn in sequons present in the same protein as sequons on the x-axis. Color is association score, proportion of proteins in which both sequons occur: brown, y-axis sequon is always present; yellow, sequons are never observed in the same protein.

### Cloning, Purification and Analysis of OsRGLP1 from S. cerevisiae

The conservation of an *N*-glycosylation site in *Os*RGLP1 suggested its importance in protein structure, stability or function. To obtain purified glycosylated protein to test the importance of its glycosylation, we expressed *Os*RGLP1 as a secreted MBP-fusion protein in *S. cerevisiae* and purified it with amylose-agarose resin. The purified protein was analyzed via SDS-PAGE and observed as a band of ~64 kDa (Fig. 2A). Top-down mass spectrometry detected four different isoforms of purified MBP-*Os*RGLP1: the most abundant form at 66,492 Da, consistent with *O*-glycosylation with 12 mannoses; 66,652 Da, with 13 *O*-mannoses; 66,815 Da, with 14 *O*-mannoses; and 68,193 Da with 12 *O*-mannoses and a GlcNAc_2_Man_8_ *N*-glycan (or a different distribution between *O*-mannoses and the length of the *N*-glycan) (Fig. 2B). These results are consistent with MBP-*Os*RGLP1 produced in *S. cerevisiae* being *O*-mannosylated at diverse sites, and partially *N*-glycosylated at a single site. To identify the specific sites of glycosylation on MBP*-Os*RGLP1 purified from *S. cerevisiae*, we performed a tryptic digest and detected glycopeptides with LC-ESI-MS/MS. The tryptic peptide containing the sole *N*-glycosylation sequon in *Os*RGLP1, V_73_GS**NVT**LINVMQIPGLNTLGISIAR_97_, was identified in unglycosylated and *N-*glycosylated high mannose forms from GlcNAc_2_Man_8_ to GlcNAc_2_Man_12_ (Supplementary Table S1, Supplementary Figure S3). Glycosylation occupancy at this site was approximately 50%, and the most abundant glycoform observed was GlcNAc_2_Man_8_. No *O*-linked glycopeptides were detected in this analysis. These results were consistent with our top-down MS analysis, and describe partial *N*-glycosylation at the single *N*-linked glycosylation sequon in MBP-*Os*RGLP1, as well as heterogeneous *O*-mannosylation across the protein.

**Figure 2.**
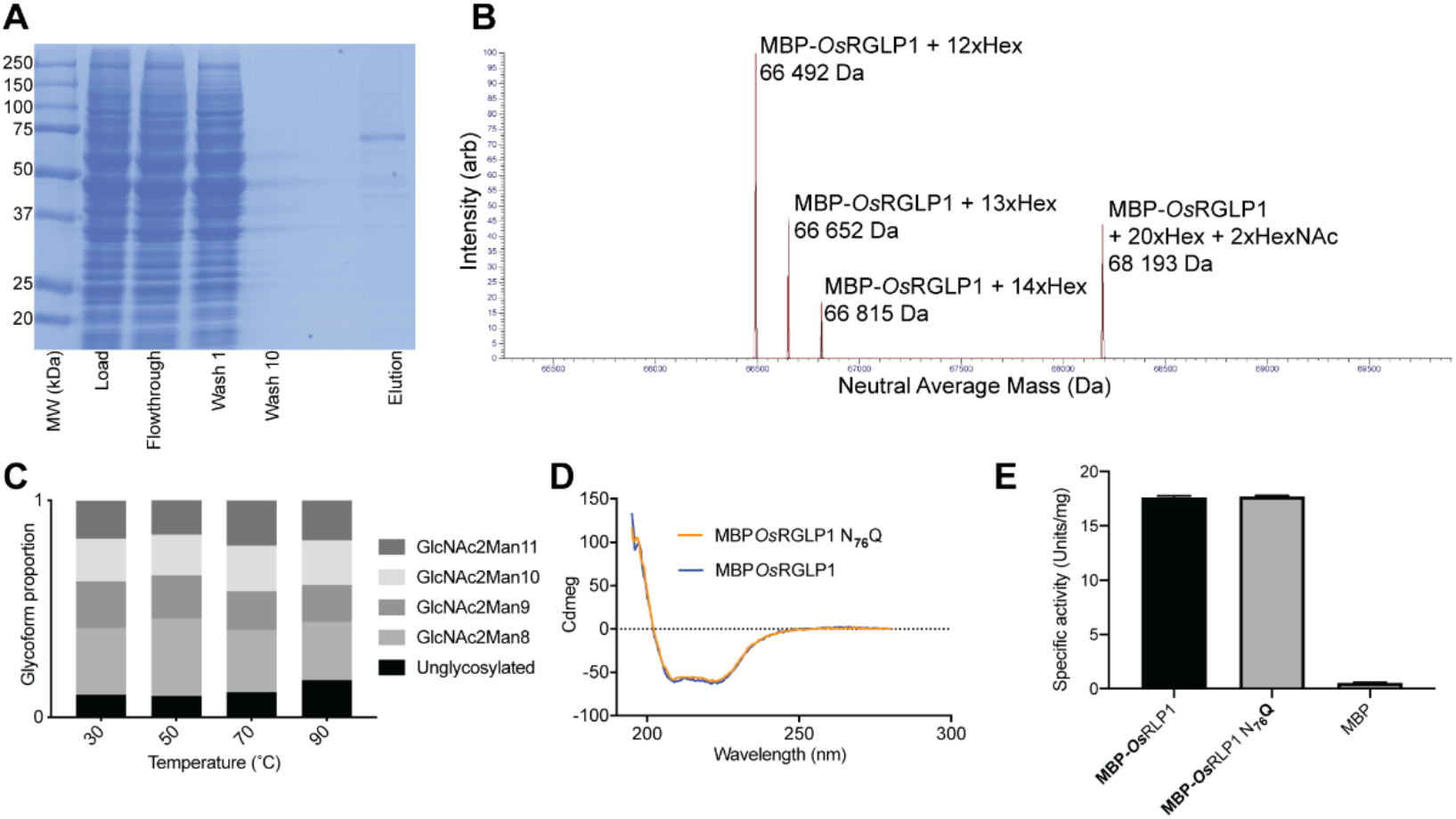
Glycosylation at the conserved *Os*RGLP1 N_76_ is not required for activity. **(A)** SDS-PAGE showing amylose-agarose resin purification of MBP-*Os*RGLP1 from *S. cerevisiae*. **(B)** Intact mass analysis of purified MBP-*Os*RGLP1. Deconvoluted spectrum showing neutral average mass. **(C)** Proportion of glycoforms at the single *N*-glycosylation site in MBP-*Os*RGLP1 in soluble protein after incubation at different temperatures as measured by LC-MS/MS. **(D)** CD spectra of MBP-*Os*RGLP1 and non-glycosylated MBP-*Os*RGLP1-N_76_Q. **(E)** In-solution activity of MBP-*Os*RGLP1, MBP-*Os*RGLP1-N_76_Q, and MBP control.

We next tested the impact of glycosylation at the single conserved *N*-glycosylation site in *Os*RGLP1 on protein stability and activity. We first performed thermal glycoprotein stability profiling of purified MBP-*Os*RGLP1, with LC-MS/MS measurement of the glycosylation profile of protein that remained soluble after incubation at temperatures from 30-90 °C (Fig. 2C). No pronounced differences were observed in the relative abundance of glycosylated and unmodified forms in the soluble fraction after incubation at any temperature. As an alternative method for measuring the impact of glycosylation on the structural integrity of *Os*RGLP1, we created a non-glycosylated MBP-*Os*RGLP1-N_76_Q variant, expressed and purified this variant and MBP-*Os*RGLP1, and measured their secondary structure by CD spectroscopy. This analysis showed equivalent CD spectra for MBP-*Os*RGLP1 and MBP-*Os*RGLP1-N_76_Q (Fig. 2D). Together, these thermal glycoprotein profiling and secondary structural analysis results showed that the presence or absence of glycosylation at the single conserved glycosylation site of MBP-*Os*RGLP1 did not have a strong influence on the thermal stability of the protein.

Overexpression of *Os*RGLP1 has been reported to increase the SOD activity of transgenic plants [26]. To test if *N*-glycosylation affected the SOD activity of MBP-*Os*RGLP1, we expressed and purified MBP-*Os*RGLP1, non-glycosylated MBP-*Os*RGLP1-N_76_Q variant, and MBP from *S. cerevisiae*, and measured their activities using an in-solution SOD assay (Fig. 2E). This assay showed clear SOD activity for both the wild-type and non-glycosylated variant proteins, which were not significantly different. This is particularly surprising, given the conservation of its single *N*-glycosylation site and highlights that evolutionary conservation does not necessarily correlate with functional importance. In summary, analysis of the expression, purification, and characterization of *Os*RGLP1 as a fusion protein from yeast showed that its single conserved *N*-glycosylation site was not required for *Os*RGLP1 stability or activity.

### Purification of OsRGLP1 from K12 Shuffle E. coli

Our analysis of MBP-*Os*RGLP1 from *S. cerevisiae* demonstrated that *N*-glycosylation was not required for *Os*RGLP1 stability or activity. However, we detected heterogeneous *O*-mannosylation of the protein, which may have complicated interpretation of these results. We therefore next attempted expression and purification of *Os*RGLP1in *E. coli*. Since *Os*RGLP1 has two cysteines that form a disulfide bond, we selected K12 shuffle *E. coli* to express and purify N-terminally His-tagged *Os*RGLP1 in an oxidized, properly folded, and active form. We could purify *Os*RGLP1 from *E. coli*, together with an abundant 60 kDa contaminant (Fig. 3A) that LC-MS/MS analysis identified as *E. coli* chaperonin GroL (Supplementary Figure 4). Expression at 16°C reduced, but did not remove, GroL contamination (Figure 3A). Interactions between molecular chaperones such as GroL and heterologously expressed proteins can result in their copurification. To remove the bacterial chaperones from *Os*RGLP1, we washed the His-*Os*RGLP1 bound talon resin with MgATP and a misfolded bacterial protein extract as described [22]. This resulted in essentially pure *Os*RGLP1 (Figure 3B). The intact mass of *Os*RGLP1 purified from *E. coli* was determined by MS to be 25,467 Da (Figure 3C), 2 Da smaller than the predicted monoisotopic mass of 25,469 Da, consistent with the presence of a disulfide bond between Cys32 and Cys47. This suggested that *Os*RGLP1 purified from K12 shuffle *E. coli* was correctly folded.

**Figure 3.**
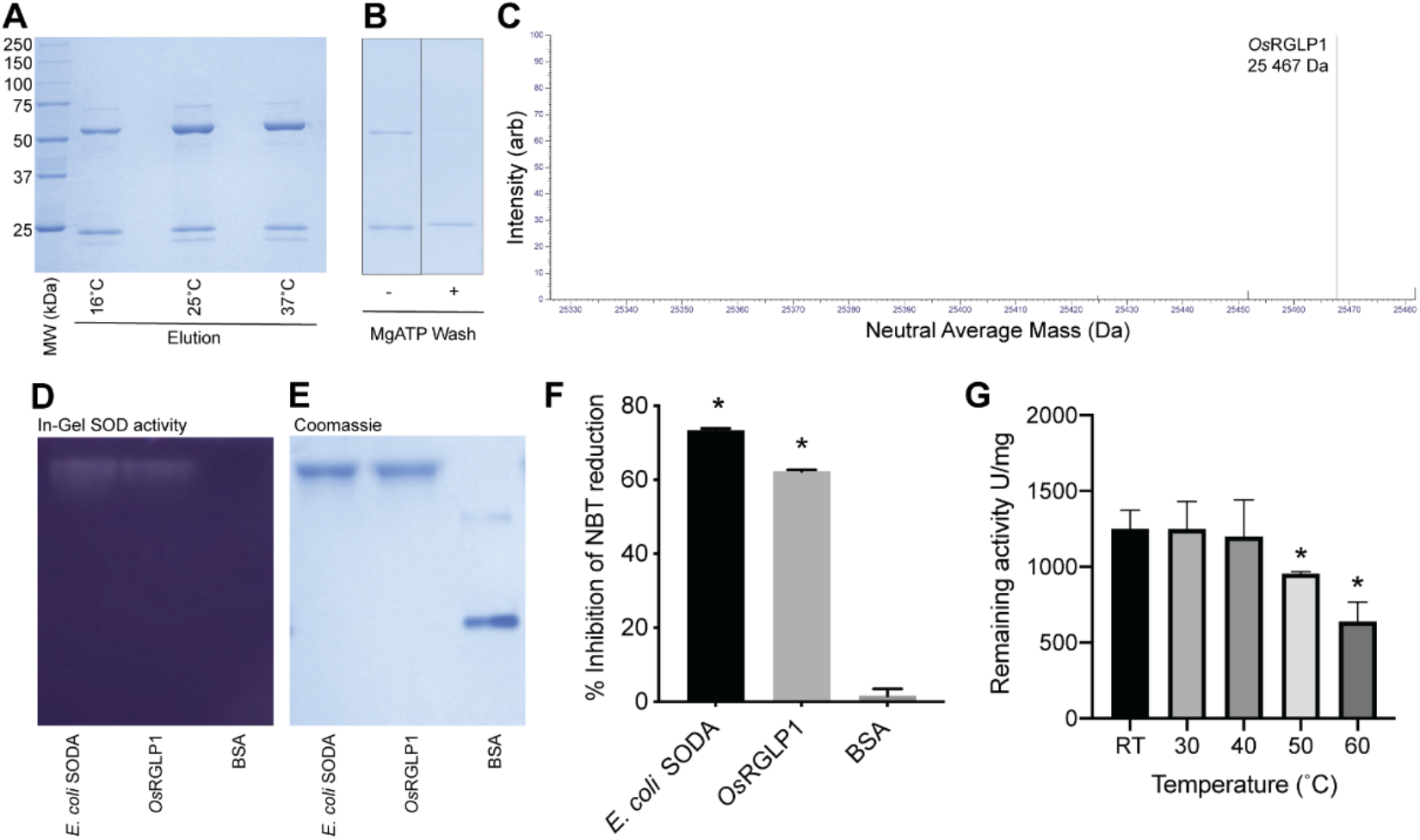
*Os*RGLP1 purified from K12 shuffle *E. coli* with MgATP wash has thermostable SOD activity. **(A)** SDS-PAGE of *Os*RGLP1 and co-purified *E. coli* GroL from K12 shuffle *E. coli* grown at different temperatures. **(B)** SDS-PAGE of *Os*RGLP1 purified without and with additional MgATP and misfolded bacterial protein extract wash. **(C)** Intact mass analysis of His-tagged *Os*RGLP1 purified from *E. coli*. Deconvoluted spectrum showing neutral average mass. **(D)** In-gel SOD activity assay (SOD activity appears as clear bands) and (**E**) replicate Coomassie stained seminative PAGE, SOD activity appears as clear bands in (D). **(F)** Quantitation of in-gel SOD activity. Values, mean. Error bars, S.D. *, *p*<0.05. **(G)** Remaining *Os*RGPL1 SOD activity after incubation at various temperatures. Values, mean. Error bars, S.D. *, *p* < 0.05 compared to RT.

We tested the SOD activity of *Os*RGLP1 purified from *E. coli* using an in-gel assay. This assay showed that *Os*RGLP1 had equivalent activity to *E. coli* SODA (Figure 3D-F), confirming that *Os*RGLP1 purified from K12 shuffle *E. coli* was properly folded and active. *Os*RGLP1 expressed in plants is reported to confer high-temperature resistant SOD activity [15, 16, 26]. To test the thermal stability of *Os*RGLP1 expressed from *E. coli*, purified protein was incubated at different temperatures and then tested for residual SOD activity (Figure 3G). No significant reduction in residual *Os*RGLP1 activity was observed after incubation at 30°C or 40°C, whereas incubation at 50°C or 60°C caused a significant reduction in residual activity. We therefore concluded that *Os*RGLP1 retained activity after incubation over the range of temperature tested, but lost activity after incubation at particularly high temperatures. The specific SOD activity of purified *Os*RGLP1 was estimated to be 1251 U/mg (Fig. 3G). This is higher than the activity for SOD from *Cicer arietinum* (157.5 U/mg) [27], but less than that of Cu-Zn SOD (SOD_C1) from citrus lemon (7456 U/mg) [28], Cu/Zn SOD from *Cucurbita moschata* (1794 U/mg) and CuZn SOD from *Allium sativum* (2867 U/mg) of SOD [29].

In summary, we successfully used two different host systems, *S. cerevisiae* and K12 shuffle *E. coli*, for expression and purification of *Os*RGLP1. This represents the first report of successful expression and purification of germins or GLPs in soluble and active form in these host systems. We provide the first direct biochemical evidence that *Os*RGLP1 has SOD activity. Finally, we found that although there was a highly conserved *N*-glycosylation sequon in *Os*RGLP1 that can be glycosylated, modification of this site does not affect protein stability, and was not required for enzymatic activity. The possibility of expressing *Os*RGLP1 in *E. coli* opens possibilities for efficient production of this protein for fundamental research and a variety of biotechnological applications.

## Acknowledgements

We thank The University of Queensland, School of Chemistry and Molecular Biosciences Mass Spectrometry Facility for assistance and expertise.

## Supplementary Material

**Supplementary Figure S1.**
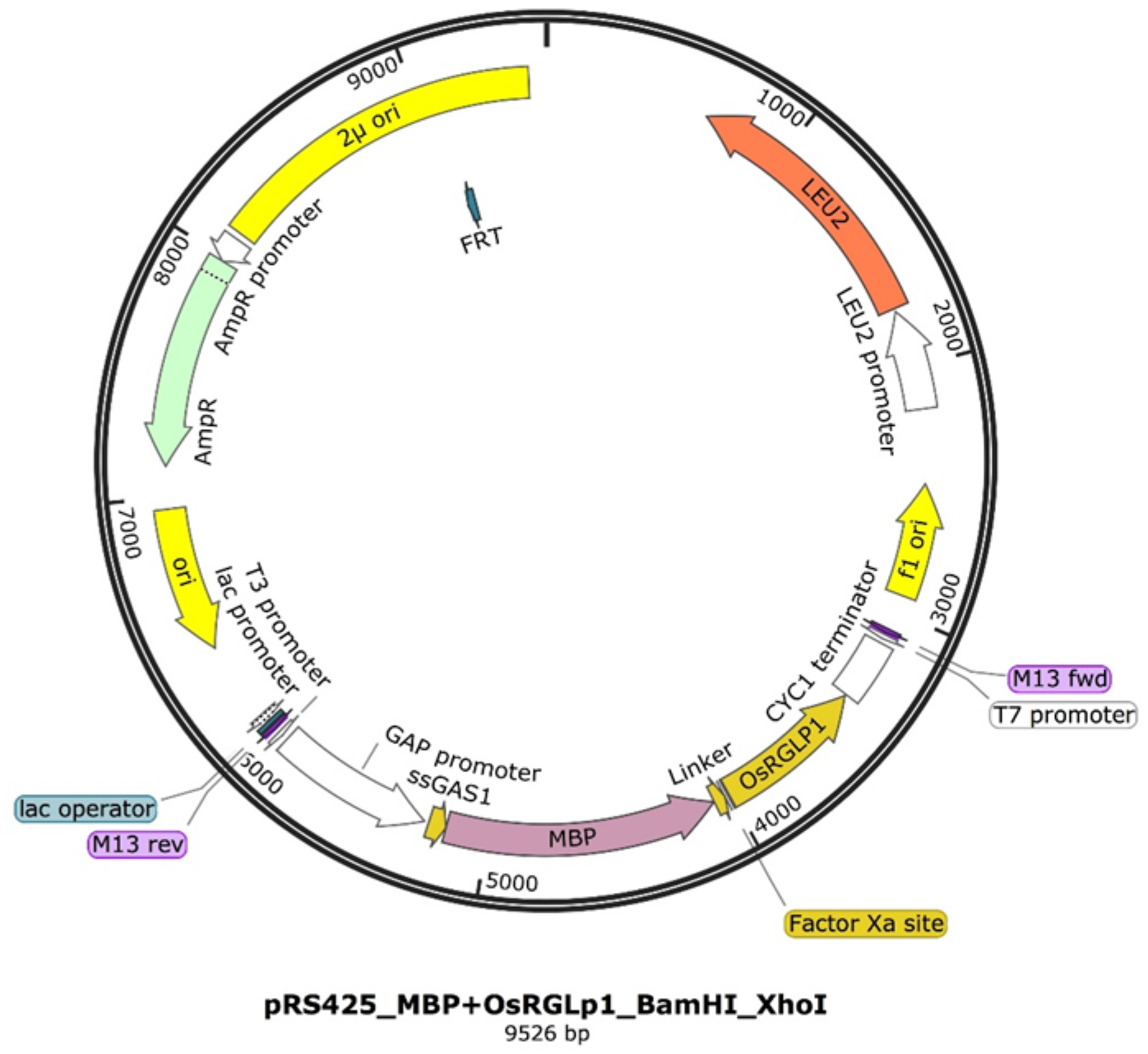
Map of pRS425-MBP-*Os*RGLP1 expression vector.

**Supplementary Figure S2.**
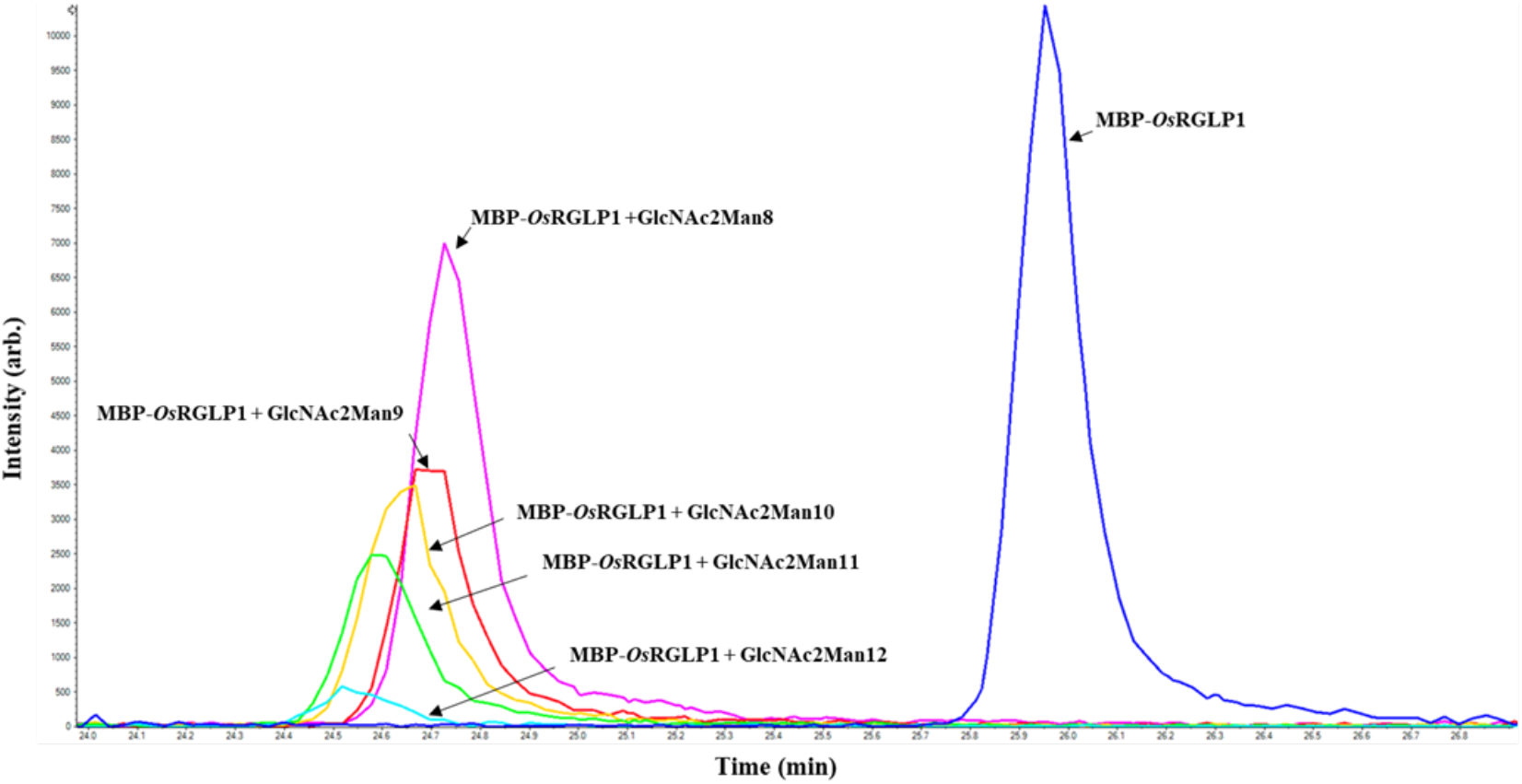
LC-MS/MS analysis of tryptic *N*-glycopeptides from MBP-*Os*RGLP1 purified from *S. cerevisiae*. Extracted ion chromatograms of glyco/peptides corresponding to V_73_GS<u>NVT</u>LINVMQIPGLNTLGISIAR_97_ unglycosylated or modified with GlcNAc_2_Man_8-12_.

**Supplementary Table S1.**
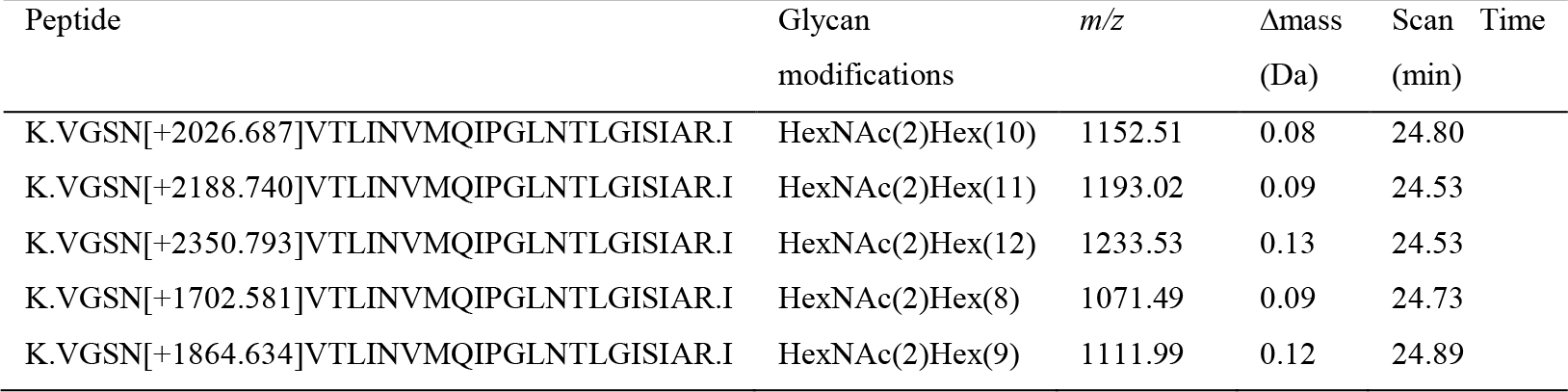
*N*-linked glycopeptides from *Os*RGLP1 identified by Byonic.

